# Pan-Peroxisome Proliferator-Activated Receptor Agonist IVA337 Alleviates Secondary Lymphedema via Inhibiting TGFβ/SMAD2/3 Signaling Pathway

**DOI:** 10.64898/2026.07.20.739660

**Authors:** Jingjing Pang, Long Nguyen Hoang Do, Esteban Delgado, Jianan Zhao, Liam Flynn, Hongxia Liu, Michael Autieri, Xiaofeng Yang, Xiaolei Liu

## Abstract

Lymphedema is a chronic disease characterized by impaired lymph drainage and accumulation of protein-rich interstitial fluid, which progresses to develop irreversible fibrosis. Importantly, effective therapies and treatments are lacking to alleviate and mitigate the disease. The pathological inflammation and fibrogenesis underlying lymphedema prompted us to evaluate a preclinical medicine IVA337, a pan-peroxisome proliferator-activated receptor (PPAR) agonist that improves liver fibrosis in patients with metabolic dysfunction-associated steatohepatitis (MASH) by activating three PPAR isoforms (α, β/δ, λ), which play critical roles in lipid metabolism, anti-inflammation responses, and anti-fibrogenesis. Here, we investigate the therapeutic effects of IVA337 during the early stage of surgery-induced secondary lymphedema in mice and explored the underlying mechanisms. IVA337 administration alleviated lymphedema progression, improved lymphatic drainage, reduced dermal thickness, and resolved lymphatic vessel dilation. Mechanistically, IVA337 suppressed the TGFβ/SMAD2/3 signaling pathway, reduced immune cells infiltration, and improves lymphatic vessels integrity. In human dermal lymphatic endothelial cells (HDLECs), IVA337 attenuated TGFβ induced SMAD2/3 phosphorylation and preserved the expression of cell junction Claudin5, reduced VE-Cadherin-stained cell-cell gaps. Collectively, our findings demonstrate that IVA337 protects against early stage lymphedema by inhibiting TGFβ/SMAD2/3-mediated inflammatory and fibrotic responses. This study provides a potential therapeutic strategy to improve lymphatic function during the early phase of lymphedema and prevent progressive fibrosis in patients with lymphedema and related disorders.

## 1. Introduction

Lymphedema is a progressive chronic lymphatic disorder with a prevalence of up to 250 million people worldwide and an estimated 10 million in the United States ^[1–3]^. Persistent lymph stasis induces chronic inflammation, immune cell infiltration, and TGFβ-mediated fibrosis, leading to irreversible tissue remodeling and lymphatic dysfunction ^[1, 4–6]^. Although compression-based therapies and surgical interventions can alleviate symptoms, no curative pharmacological treatment is currently available ^[7,8]^. Therefore, targeting early inflammatory and fibrotic responses represents a promising strategy to preserve lymphatic function and prevent disease progression.

PPARs are ligand-activated nuclear receptors that regulate transcriptional programs involved in lipid metabolism, inflammation, immune responses, vascular biology, and tissue remodeling ^[9–11]^. Based on these effects, PPAR agonists such as fibrates (PPARα agonist) and thiazolidinediones (PPARγ agonist) have been clinically used mainly for hypertriglyceridemia and insulin resistance ^[12, 13]^. In particular, a new pan-PPAR agonist IVA337 (also known as lanifibranor) can simultaneously activate three PPAR isoforms; this balanced activation profile allows it to modulate multiple biological pathways involved in lipid and glucose metabolism, inflammation, and fibrosis. Many studies report that IVA337 treatment could alleviated liver fibrosis in MASH and skin fibrosis ^[14-16]^. PPARγ activation has been shown to reduce fibroadipose remodeling in secondary lymphedema by regulating PDGFRα^⁺^ fibroadipogenic mesenchymal cells ^[17]^, and activation of PPAR signaling can also suppresses inflammatory and TGFβ-driven fibrotic responses ^[18–21]^, therefore we hypothesized that IVA337 could attenuate early-stage secondary lymphedema by suppressing inflammatory and TGFβ-mediated fibrotic responses while preserving lymphatic endothelial junction integrity.

In this study, we demonstrate that IVA337 alleviates tail surgery-induced secondary lymphedema by inhibiting TGFβ/SMAD2/3 signaling pathway. IVA337 treatment reduces dermal thickening and lymphatic vessel dilation, improves lymphatic drainage function, restores lymphatic vessel integrity, and decreases inflammatory cells infiltration. Furthermore, in primary HDLECs, IVA337 pretreatment attenuates TGFβ-induced SMAD2/3 phosphorylation, preserves Claudin5 (tight junction protein) expression, and reduces intercellular gap formation. Collectively, these findings identify a previously unrecognized function of IVA337 in the regulation of lymphatic function and provide new insights into potential therapeutic strategies for lymphedema.

## 2. Materials and methods

### 2.1. Animals and Treatments

All mice used in this study were maintained on C57BL/6J genetic background. 8 weeks old mice were used for the surgery-induced secondary lymphedema, male and female were included. In brief, Mice were anesthetized with isoflurane throughout the procedure. Tail diameter was measured at 5 mm intervals starting 20 mm from the tail base prior to surgery. A 3 mm circumferential full-thickness skin excision was performed 20 mm from the tail base. Postoperatively, mice were monitored daily for 3 weeks and tail volumes were quantified after the surgery at day 0, 3, 7, 14, and 21. IVA337 (Selleck Chemicals,S8770) was administered 20 mg/kg by intraperitoneally injection after the surgery at day 0,3, 7,10, 14, and 17. Lymphatic drainage function was evaluated by FITC-dextran (70kDa, Sigma-Aldrich,46945) tracer injection and tail picture was captured after the injection at 2h and 4h, the tracer clearance rate was quantified to evaluate lymphatic drainage function. All animal experiments and husbandry procedures were performed in accordance with protocols approved by the Temple University Institutional Animal Care and Use Committee.

### 2.2 Cell culture and Treatments

Human dermal LECs were purchased from Promocell (C-22121) and cultured with endothelial basal medium complemented with supplement mix (Promocell, C-39221) and 5% penicillin/streptomycin. Experiments were performed using P4-P7 passages. HDLECs were pretreated with IVA337 (5 µM, 16h) and then treated with TGFβ (20 ng/ml, 30 min; Thermofisher, 100-35B) to induce SMAD phosphorylation or TGFβ (20 ng/ml, 8h) to check cell junctional protein.

### 2.3 Immunofluorescence Staining and Confocal Microscopy

For whole-mount immunostaining, ear tissues were fixed in 2% paraformaldehyde (PFA) overnight at 4°C. Fixed tissues were washed with phosphate-buffered saline (PBS) and blocked in blocking buffer overnight at 4°C. Samples were then incubated with primary antibodies overnight at 4°C, followed by washing with PBST (PBS containing 0.1% Triton X-100). Appropriate secondary antibodies were applied for 2 hours at room temperature, after which tissues were washed with PBST and mounted using ProLong mounting medium (Sigma-Aldrich, P36931).

For cell immunostaining, cells were fixed with 2% PFA at room temperature and washed four times with PBS. Samples were then blocked for 1 hour in blocking solution, followed by incubation with primary antibodies for at least 2 hours at room temperature. Cells were subsequently washed with PBST and incubated with fluorophore-conjugated secondary antibodies diluted in blocking solution for 2 hours. After additional washes with PBST, samples were mounted using ProLong mounting medium with DAPI (Cell Signaling Technology, 8961S).

Confocal images were acquired using a Nikon AX laser scanning confocal microscope mounted on an Eclipse Ti2 inverted microscope stand (Nikon Instruments). The system was equipped with a DUX-ST detector unit and LUD-S4 laser light source, and images were collected using NIS-Elements software (Nikon).

Detailed information on all primary and secondary antibodies used in this study is provided in Supplementary Table S1.

### 2.4 Hematoxylin and eosin (H&E) staining

Cryosections (10 μm) of the tail were fixed in 2% paraformaldehyde for 10 min, rinsed with phosphate-buffered saline (PBS), and stained with hematoxylin followed by eosin using standard procedures. Briefly, sections were immersed in hematoxylin, differentiated, blued, and counterstained with eosin. After dehydration through graded ethanol, clearing in xylene, and mounting with a permanent mounting medium, stained sections were imaged using a bright-field microscope.

### 2.5 Western Blot

To examine signaling changes in response to various treatments, cells were lysed in RIPA buffer (Invitrogen, PI89900) supplemented with a protease inhibitor cocktail (Pierce, A32959). Protein concentrations were determined using the BCA Protein Assay Kit (Thermo Fisher Scientific, 23227). Equal amounts of protein were subjected to SDS– PAGE, transferred to PVDF membranes, and analyzed by Western blotting. Protein bands were detected using SuperSignal™ West Pico Plus chemiluminescent substrate (Thermo Fisher Scientific) and imaged with the iBright™ 1500 Imaging System (Thermo Fisher Scientific). Band intensities were quantified using ImageJ (version 2.14.0). All raw data used for quantification are provided in the Supplementary Information, and details of primary and secondary antibodies are listed in Supplementary Table S1.

### 2.6 Quantitative Real-Time PCR (qRT–PCR)

Total RNA was extracted from mouse tail or HDLECs using TRIzol reagent (Thermo Fisher Scientific, 15596018) according to the manufacturer’s instructions. RNA concentration and purity were determined prior to downstream analysis. Complementary DNA (cDNA) was synthesized from total RNA using the iScript cDNA Synthesis Kit (Bio-Rad, 1708891). Quantitative real-time PCR (qRT–PCR) was performed using Power SYBR Green PCR Master Mix (Applied Biosystems/Life Technologies) on a StepOnePlus Real-Time PCR System (Applied Biosystems). Each sample was analyzed individually in each qPCR run, and reactions were performed in technical replicates. Gene expression levels were normalized to the housekeeping gene GAPDH, and relative expression was calculated using the 2^−ΔΔ^Ct method. Primer sequences used in this study are listed in Supplementary Table S2.

### 2.7 Statistics

All data are presented as mean ± standard error of the mean (SEM) unless otherwise indicated. Statistical analyses were performed using GraphPad Prism. Comparisons between two groups were conducted using an unpaired two-tailed Student’s t-test, whereas comparisons among multiple groups were analyzed using one-way ANOVA followed by Turkey’s test. The number of biological replicates (n) is indicated in the corresponding figure legends. Differences were considered statistically significant when P < 0.05. All experiments were repeated independently at least three times unless otherwise stated.

### 2.8 Study Approval

All animal experiments were performed in accordance with the guidelines of the Institutional Animal Care and Use Committee (IACUC) of Temple University. All experimental protocols were reviewed and approved by the Temple University IACUC, and all procedures were conducted in compliance with the Guide for the Care and Use of Laboratory Animals published by the National Institutes of Health (NIH).

## 3. Results

### 3.1 IVA337 attenuates tail surgery-induced secondary lymphedema

To investigate the therapeutic effect of IVA337 on secondary lymphedema, we established a mouse tail surgery-induced lymphedema model by disrupting tail lymphatic vessels (Fig. 1A). The monitor time point and IVA337 injection time point were as depicted in S-Fig. 1A. To assess the potential systemic effects of IVA337 administration, we monitored mouse body weight throughout the experimental period. No significant differences in body weight between vehicle- and IVA337-treated groups were observed after tail surgery (S-Fig. 1B), suggesting that IVA337 administration was not associated with systemic changes. To confirm PPAR pathway activation following IVA337 treatment, we isolated RNA from tail skin tissues and examined the expression of canonical target genes for PPARs, including *Cpt1a* and *Fabp4*^[22]^. IVA337 treatment significantly increased the expression of *Cpt1a* and *Fabp4* (S-Fig. 2A–B). These results indicate that IVA337 activates PPAR signaling in skin tissues. Compared with vehicle-treated controls, IVA337 administration markedly reduced tail swelling at day 21 after surgery (Fig. 1B). Longitudinal measurement of tail volume at days 3, 7, 14, and 21 demonstrated that IVA337 significantly attenuated postoperative tail edema progression (Fig. 1C). Histological analysis by H&E staining revealed that IVA337 treatment reduced epidermal and dermal/hypodermal thickening compared with the vehicle group (Fig. 1D–F), indicating decreased tissue remodeling during lymphedema development. To evaluate lymphatic drainage function, 10 ul FITC-dextran was injected into the tail, and tracer clearance was assessed at 2 and 4 h after injection in mice at day 21 after surgery (Fig. 1G). IVA337-treated mice exhibited significantly enhanced FITC-dextran clearance compared with vehicle-treated controls, suggesting improved lymphatic transport capacity (Fig. 1H). Because lymphatic vessel remodeling, including lymphangiogenesis and vessel dilation, contributes to lymphatic dysfunction during lymphedema progression, we performed immunostaining for lymphatic endothelial markers Lyve1 and Prox1 to assess lymphatic vessel density and morphology (Fig. 1I). IVA337 treatment significantly reduced lymphatic vessel dilation but did not alter lymphatic vessel density compared with vehicle-treated mice (Fig. 1J–K). Collectively, these findings demonstrate that IVA337 alleviates early-stage secondary lymphedema by reducing tissue swelling and dermal thickening, improving lymphatic drainage, and preserving lymphatic vessel morphology.

**Figure 1.**
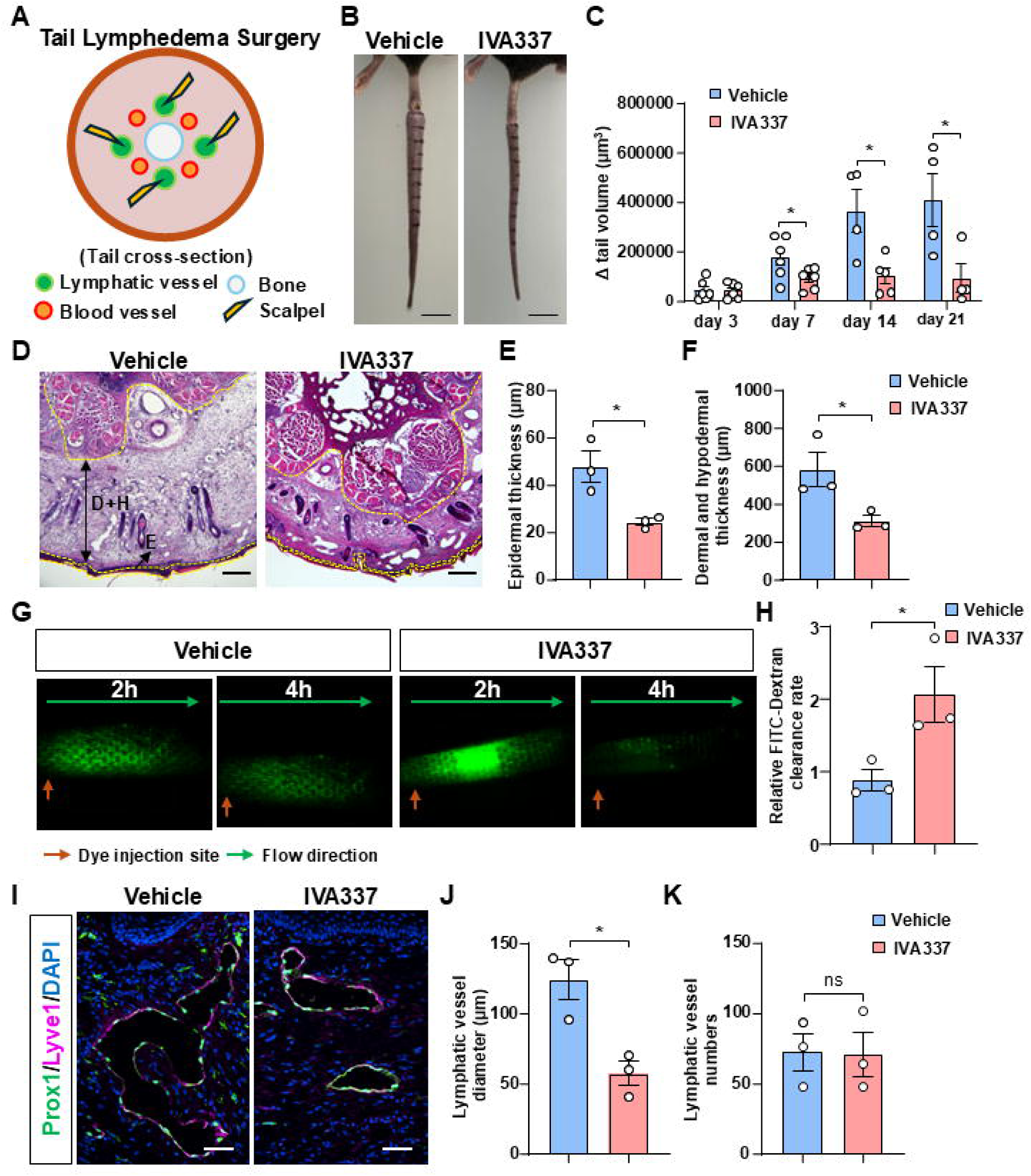
IVA337 alleviates tail lymphedema after tail lymphedema surgery. **A.** Diagram of tail lymphedema surgery model. Lymphatic vessels from tail were damaged by scalpel. **B-C.** tails were captured at 21 days; tail volume was analyzed at day 3, day 7, day 14, and day 21 after the surgery. N=4-6. **D.** H&E staining of tail cross-section at day 21 after surgery. Yellow dash line separate Epidermis (E), dermis (D), and hypodermis (H), black arrows indicate the area. **E.** Epidermis thickness was quantified and analyzed. N=3. **F.** Dermis and hypodermis thickness were quantified and analyzed. N=3. **G.** Lymphatic drainage was evaluated by the clearance of FITC-Dextran. 10 µl FITC-Dextran (1 mg/ml) was injected into the tail of day21 surgery mouse, and calculated fluorescent intensity after 2 h and 4 h. **H.** FITC-Dextran clearance rate was quantified per hour. N=3. **I.** Immunofluorescence staining for Prox1 and Lyve1 to check positive vessels of tail cryosections from day 21 surgery mice. **J-K.** Lymphatic vessel numbers and diameter were quantified for total field of tail section. N=3. Scale bars: B is 1 cm; D is 200 µm; I is 50 µm. Data are shown as mean ± SEM, *P<0.05, unpaired two-tailed Student’s t-test.

### 3.2 IVA337 inhibits SMAD2/3 phosphorylation in tail skin and tail lymphatic vessels

Given that IVA337 exhibits anti-fibrotic effects in liver fibrosis ^[15]^, we investigated whether its protective effects against secondary lymphedema are mediated through suppression of fibrosis-associated signaling pathways. As TGFβ/SMAD signaling is a central driver of tissue fibrosis, we examined SMAD2/3 phosphorylation in tail tissues following IVA337 treatment. Immunostaining of tail cryosections revealed that IVA337 administration significantly reduced the overall number of p-SMAD2/3-positive cells compared with vehicle-treated controls (Fig. 2A–B). To determine whether IVA337 directly affects LECs signaling, we further analyzed p-SMAD2/3 activation in Lyve1^⁺^/Prox1^⁺^ lymphatic endothelial cells. IVA337 treatment markedly decreased the number of Lyve1^⁺^/Prox1^⁺^/p-SMAD2/3^⁺^ cells, indicating reduced SMAD2/3 activation in lymphatic endothelial cells (Fig. 2C–D). Consistently, qRT-PCR analysis demonstrated decreased expression of the fibrosis-associated gene *Col1a1* in IVA337-treated tail tissues (Fig. 2E). Together, these findings demonstrate that IVA337 suppresses SMAD2/3 activation in both skin tissues and lymphatic endothelial cells, suggesting that inhibition of TGFβ/SMAD-mediated fibrotic signaling contributes to its protective effects against lymphedema.

**Figure 2.**
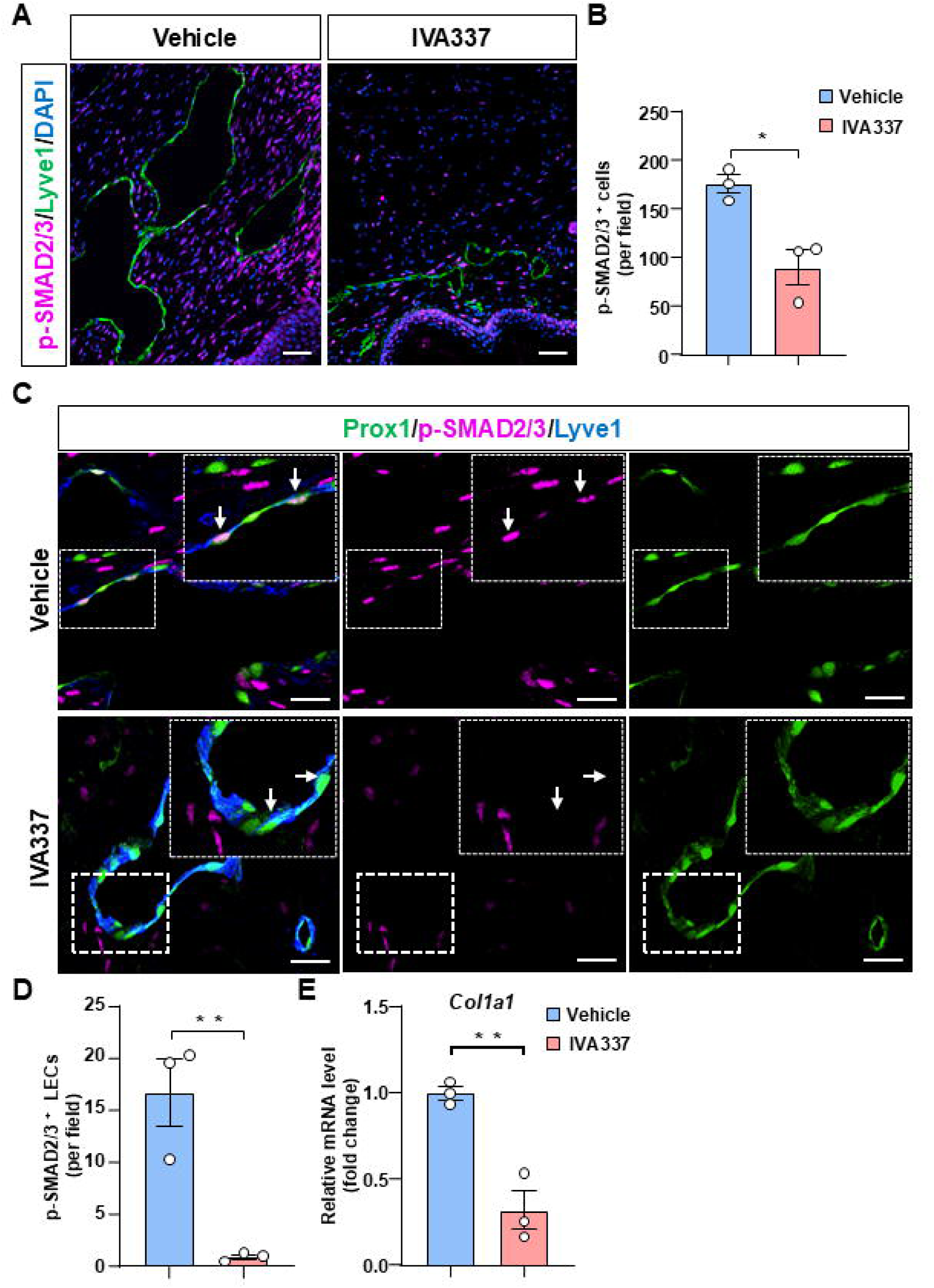
IVA337 suppresses SMAD2/3 phosphorylation in the tail skin and lymphatic vessels of secondary lymphedema mice. **A-B.** Immunostaining of tail cryosection targeted for p-SMAD2/3 and Lyve1, and quantification of p-SMAD2/3 cell numbers per field. N=3. **C-D.** Immunostaining of tail cryosection targeted for p-SMAD2/3, Prox1, and Lyve1, white arrows indicate p-SMAD2/3 positive LECs, and quantification of p-SMAD2/3 cell numbers in lymphatic endothelial cells, N=3. **E.** qRT-PCR to check fibrosis related gene *Col1a1* level in mouse tail. Scale bars: A is 50 µm; C is 20 µm. Data are shown as mean ± SEM, *P<0.05, **P<0.01, unpaired two-tailed Student’s t-test.

### 3.3 IVA337 reduces inflammatory cell infiltration in tail skin

Chronic inflammation is a critical driver of secondary lymphedema progression and contributes to lymphatic dysfunction and tissue fibrosis ^[2, 23]^. Interestingly, TGFβ/SMAD signaling is a master regulator of immune and inflammatory responses. It maintains tolerance under normal conditions but, when dysregulated, can drive chronic inflammation, fibrosis, and immune imbalance ^[24, 25]^. Based on our previous results and the proved anti-inflammatory effects of PPAR activation, we investigated whether IVA337 modulates inflammatory responses in the lymphedematous skin.

Immunostaining of tail cryosections demonstrated a reduction in CD45^⁺^ leukocytes, CD68^⁺^ macrophages, and CD8a^⁺^ cytotoxic T cells in the IVA337-treated group (Fig. 3A-F), whereas CD19^⁺^ B cells and Ly6G^⁺^ neutrophils were not significantly altered (S-Fig. 3A-D). Together, these findings suggest that IVA337 attenuates specific inflammatory cell infiltration and may contribute to improved lymphedema outcomes through anti-inflammatory mechanisms.

**Figure 3.**
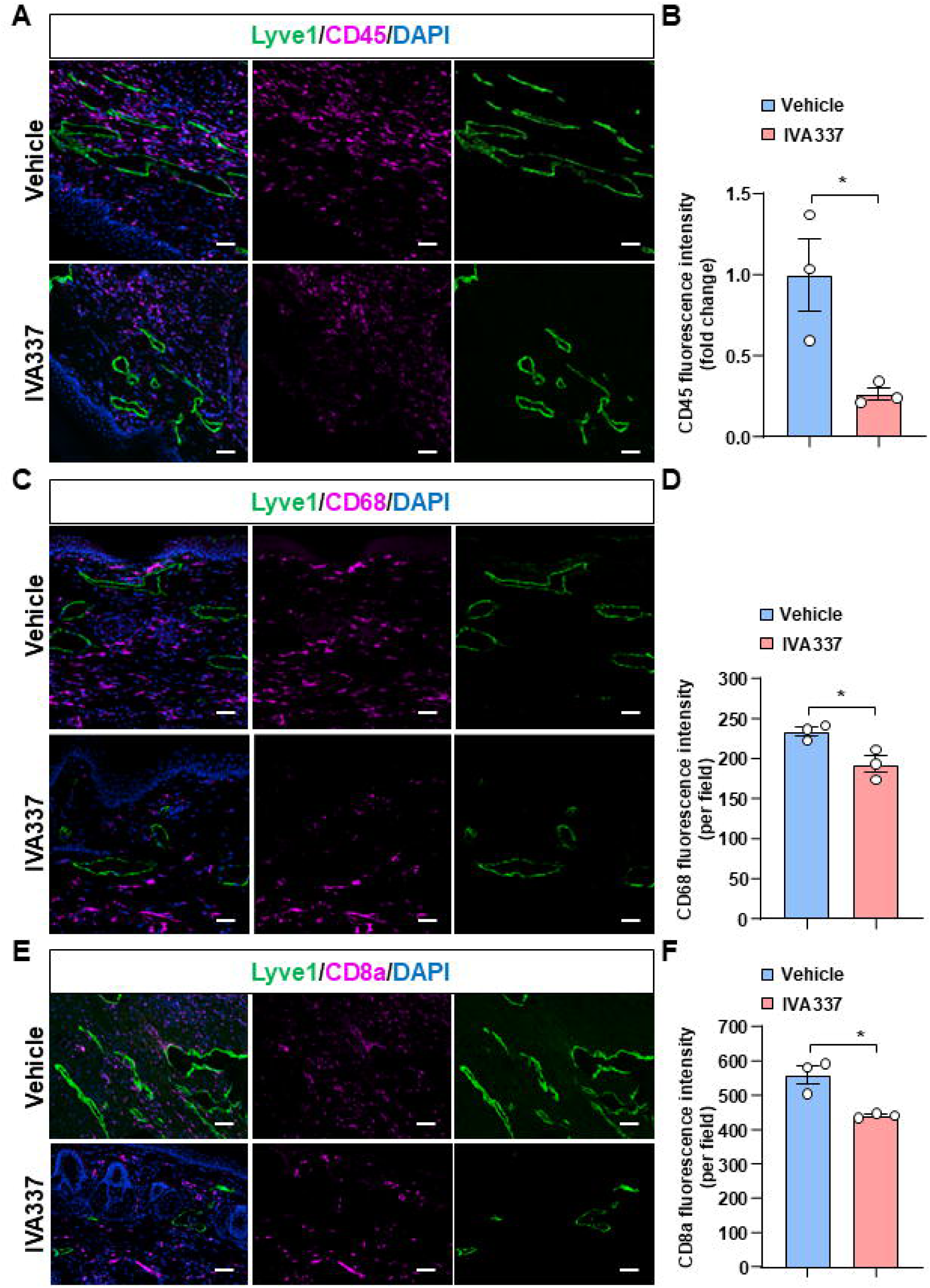
IVA337 decreases immune cell infiltration in mouse tail of lymphedema. **A-B.** Immunostaining targeted for leukocytes marker CD45 together with LECs marker Lyve1 in tail cryosection, and quantification and analysis of relative fluorescent intensity in vehicle and IVA337 group. N=3. **C-D.** Immunostaining targeted for macrophage marker CD68 and LECs marker Lyve1 in tail cryosection, and quantification and analysis of fluorescent intensity. N=3. **E-F.** Immunostaining targeted for T cell marker CD8a and LECs marker Lyve1 in tail cryosection, and quantification and analysis of fluorescent intensity. N=3. Scale bar is 50 µm. Data are shown as mean ± SEM, *P<0.05, **P<0.01, unpaired two-tailed Student’s t-test.

### 3.4 IVA337 preserves lymphatic vessel integrity by restoring junctional protein expression and reducing lymphatic gaps

Lymphatic endothelial cell junctions are essential for maintaining lymphatic vessel structure and barrier function, and disruption of these junctions contributes to lymphatic dysfunction during lymphedema early stage and its progression ^[26, 27]^. To determine whether IVA337 regulates lymphatic endothelial junction integrity, we examined the expression of tight junction proteins in tail cryosections. Immunostaining analysis revealed that IVA337 treatment restored the expression of the tight junctional proteins Claudin5 and ZO1 in lymphedematous tissues compared with vehicle-treated controls (Fig. 4A–D). Because lymphatic vessels exhibited increased structural disruption following tail surgery, we further assessed lymphatic vessel integrity by analyzing VE-Cadherin-positive intercellular gaps in Lyve1^⁺^ lymphatic vessels. Compared with vehicle-treated mice, IVA337 administration significantly reduced VE-cadherin-associated gap formation in lymphatic vessels (Fig. 4E–F). Together, these findings demonstrate that IVA337 preserves lymphatic vessel integrity during secondary lymphedema progression by restoring lymphatic endothelial junctional protein expression and reducing junctional disruption.

**Figure 4.**
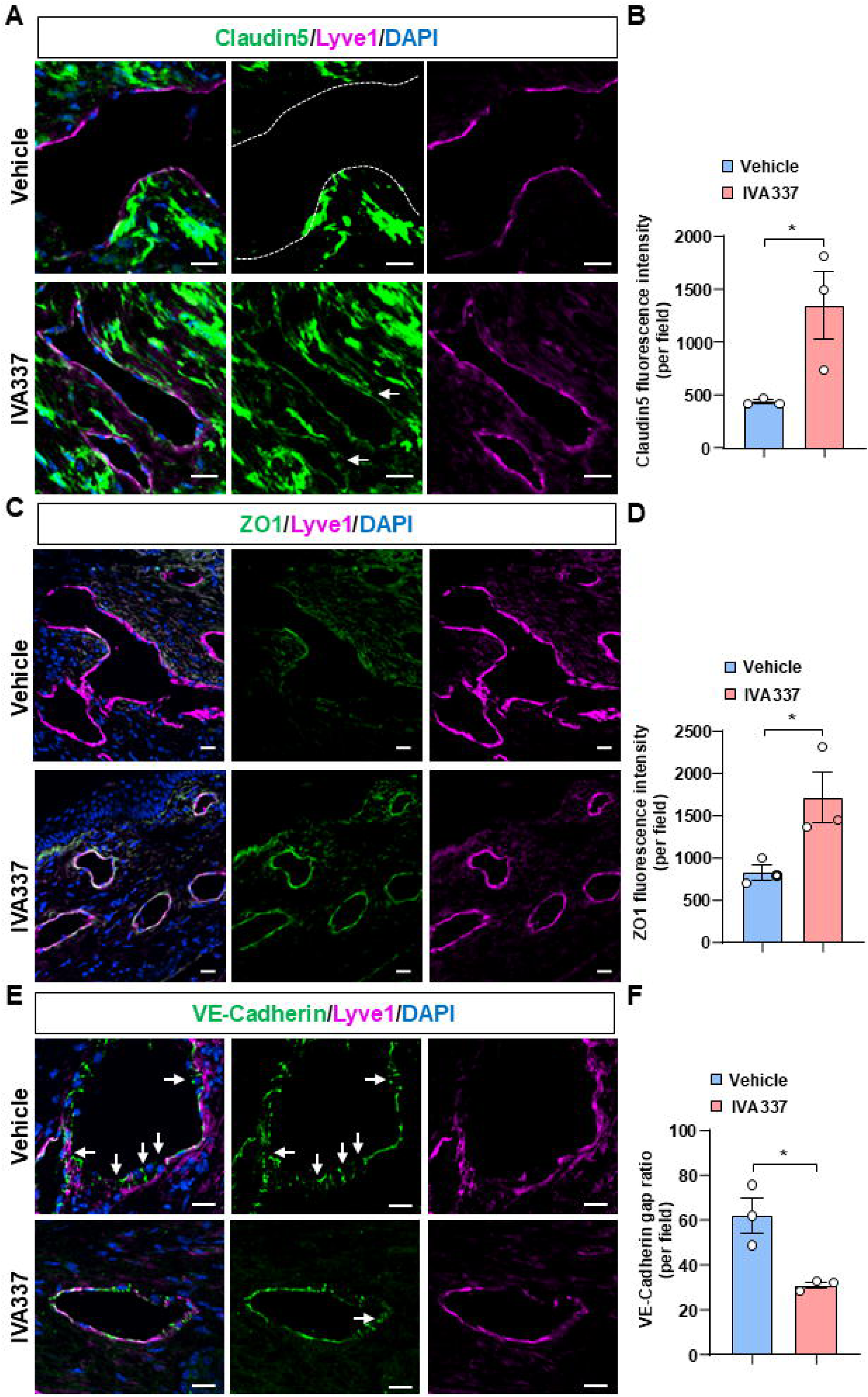
IVA337 rescues lymphatic tight junction and decreases intercellular gaps in tail lymphedema. **A-B.** Immunostaining targeted for tight junction marker Claudin5 together with LECs marker Lyve1 in tail cryosection, white dash line indicates lymphatic vessel, and quantification and analysis of relative fluorescent intensity in vehicle and IVA337 group. N=3. **C-D.** Immunostaining targeted for tight junction marker ZO1 together with LECs marker Lyve1 in tail cryosection, and quantification and analysis of relative fluorescent intensity in vehicle and IVA337 group. N=3. **E-F.** Immunostaining targeted for VE-Cadherin to label intercellular gaps together with LECs marker Lyve1 in tail cryosection, white arrows indicate intercellular gaps, and they were quantified and analyzed in vehicle and IVA337 group. N=3. Scale bar is 20 µm. Data are shown as mean ± SEM, *P<0.05, unpaired two-tailed Student’s t-test.

### 3.5 IVA337 protects lymphatic endothelial junction integrity against TGF**β**-induced dysfunction

TGFβ is a critical mediator of fibrotic progression in lymphedema ^[28]^. To determine whether IVA337 directly regulates TGFβ-induced SMAD2/3 signaling in lymphatic endothelial cells, HDLECs were pretreated with IVA337, followed by stimulation with TGFβ. Western blot analysis demonstrated that IVA337 significantly attenuated TGFβ-induced SMAD2/3 phosphorylation (Fig. 5A-B), indicating inhibition of TGFβ signaling activation. Because our in vivo findings showed that increased p-SMAD2/3 activation in lymphatic vessels was associated with reduced Claudin5 and ZO1 expression and increased VE-cadherin-positive gaps, we further investigated whether IVA337 protects lymphatic endothelial junction integrity under prolonged TGFβ stimulation. HDLECs were pretreated with IVA337 and subsequently exposed to TGFβ. TGFβ stimulation significantly reduced Claudin5 expression, whereas IVA337 pretreatment restored Claudin5 levels (Fig. 5C-D). Meanwhile, ZO1 expression was not significantly affected by TGFβ stimulation compared with controls (S-Fig 4A-B). However, IVA337 improves ZO1 expression when stimulated with TGFβ. Furthermore, TGFβ treatment increased VE-cadherin-positive intercellular gap formation, while IVA337 pretreatment markedly reduced TGFβ-induced gap formation and preserved endothelial junction organization (Fig. 5E-F). We also explored mRNA levels of *CLDN5* (Claudin5), *CDH5* (VE-Cadherin), and *TJP1* (ZO1) by qRT-PCR, and found that only *TJP1* was slightly increased in TGFβ-treated group (S-Fig 5A-C). This indicates that IVA337 restores cell junction expression is likely independent of transcriptional regulation. Collectively, these results demonstrate that IVA337 suppresses TGFβ/SMAD signaling activation and protects lymphatic endothelial integrity by maintaining Claudin5 and ZO1 expression, and reducing junctional disruption.

**Figure 5.**
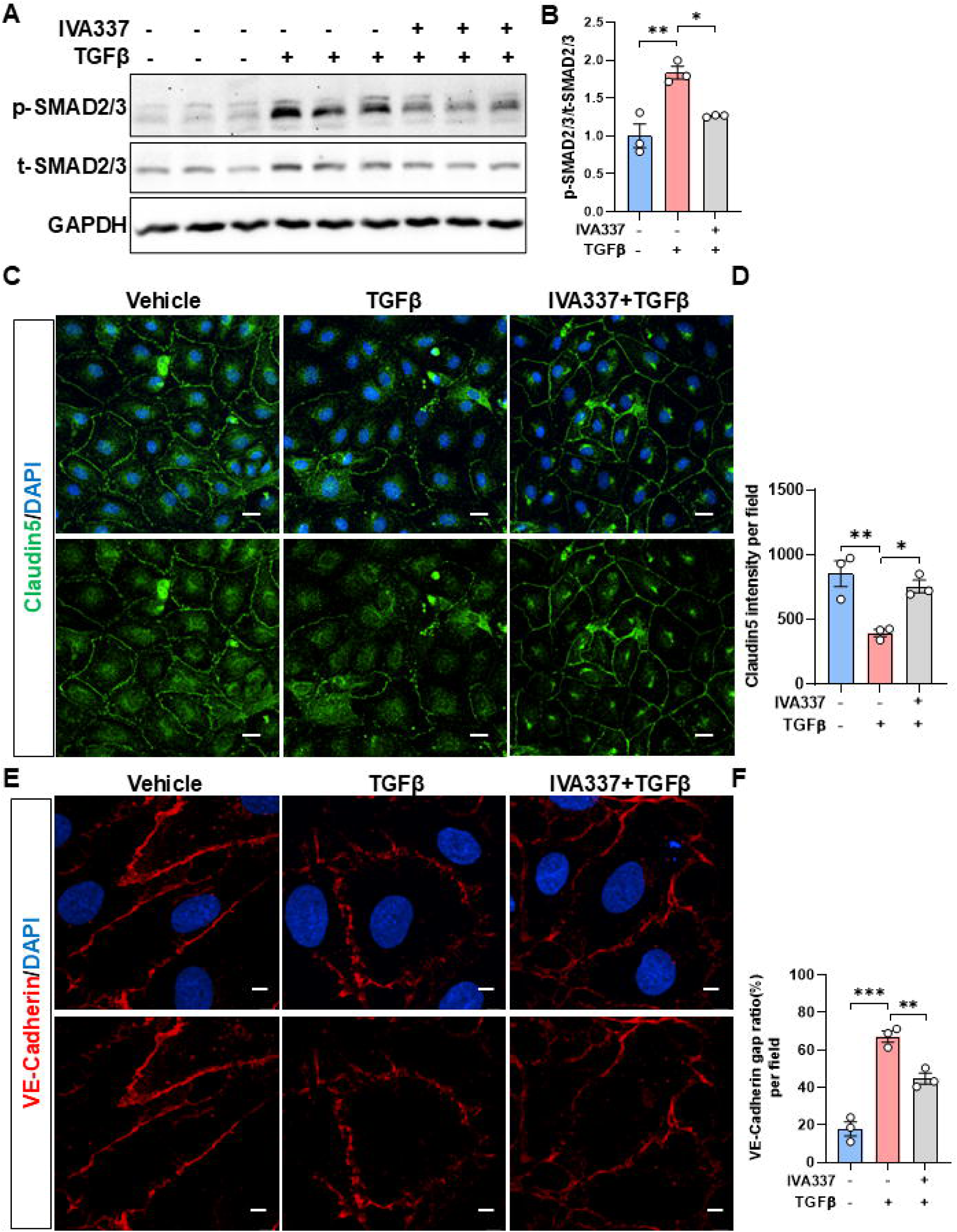
IVA337 inhibits TGFβ-induced SMAD phosphorylation and protects cell integrity. **A-B.** HDLECs were pretreated IVA337 (5 µM, 16h) and then treated with TGFβ (100 ng/ml, 30 min), western blot targeted for p-SMAD2/3, t-SMAD2/3, and GAPDH to check the protein levels. Quantification and analysis were shown in B. N=3. **C-D.** HDLECs were pretreated IVA337 (5 µM, 16h) and then treated with TGFβ (20 ng/ml, 8 h), immunostaining targeted for Claudin5 and DAPI to check the cell junction. Fluorescent intensity was analyzed in D. N=3. **E-F.** HDLECs were pretreated IVA337 (5 µM, 16h) and then treated with TGFβ (20 ng/ml, 8 h), immunostaining targeted for VE-Cadherin and DAPI to check the intercellular gaps. Cellular gaps were analyzed in F. N=3. Scale bar: C is 20 µm, E is 5 µm. Data are shown as mean ± SEM, *P<0.05, **P<0.01, ***P<0.001, one-way ANOVA followed by Turkey’s test.

In conclusion, our findings demonstrate that IVA337 alleviates secondary lymphedema by suppressing TGFβ/SMAD signaling, thereby reducing dermal fibrosis and lymphatic vessel dilation, improving lymphatic drainage, preserving lymphatic endothelial junction integrity, and limiting inflammatory cell infiltration (Fig. 6).

**Figure 6.**
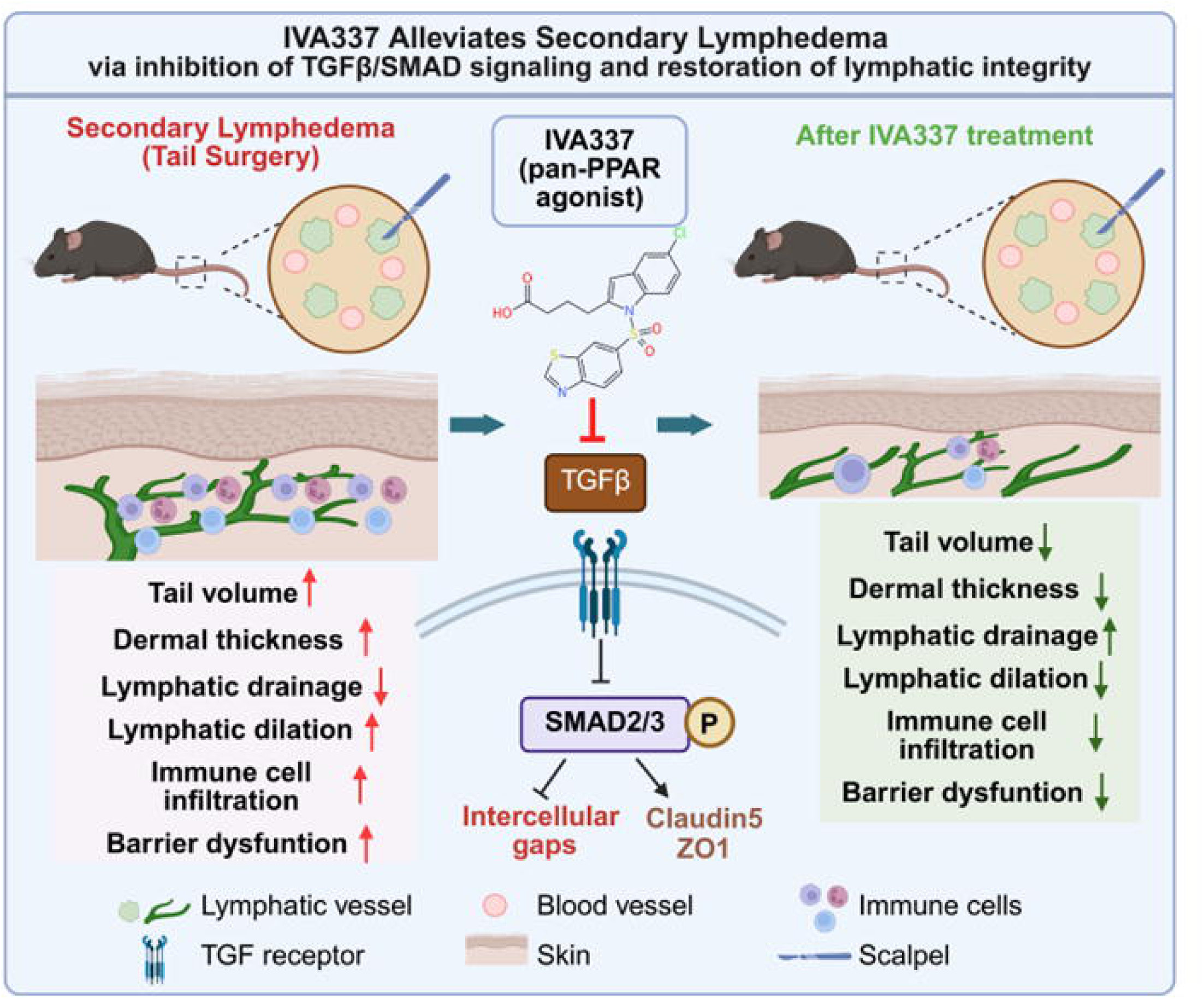
IVA337 improves secondary lymphedema via inhibiting TGFβ/SMAD signaling pathway and restores lymphatic integrity. Schematic illustration shows that IVA337 reduces dermal fibrosis and lymphatic vessel dilation, improves lymphatic drainage, preserves lymphatic endothelial junction integrity, limits inflammatory cell infiltration, and suppresses TGFβ/SMAD signaling and TGFβ-induced barrier dysfunction, all of these finally leading to the alleviation of secondary lymphedema.

## Discussion

The development of pharmacological therapies that directly target the molecular mechanisms underlying lymphatic dysfunction and fibrotic remodeling remains a major unmet need in the treatment of lymphedema ^[29]^. In this study, we identify IVA337, a pan-PPAR agonist, as a potential therapeutic strategy for the treatment of secondary lymphedema. We demonstrate that IVA337 treatment reduces tail swelling, improves lymphatic drainage, decreases dermal thickening and lymphatic vessel dilation, and preserves lymphatic endothelial integrity in a mouse tail surgery-induced lymphedema model. Mechanistically, IVA337 suppresses TGFβ/SMAD2/3 signaling, reduces inflammatory cell infiltration, and protects lymphatic endothelial junctions.

Following lymphatic injury, protein-rich lymph accumulation promotes immune cell recruitment and inflammatory cytokine production, which further activates fibroblasts and drives extracellular matrix deposition ^[30]^. Among these pathways, TGFβ signaling is a central regulator of fibrosis by inducing SMAD2/3 phosphorylation and promoting myofibroblast activation ^[31]^. Previous studies have demonstrated that inhibition of TGFβ signaling reduces fibrosis and improves lymphatic function in experimental lymphedema models ^[28]^. Consistent with these findings, we observed increased p-SMAD2/3 activation in lymphedematous tissues and lymphatic endothelial cells, whereas IVA337 treatment markedly reduced SMAD2/3 phosphorylation and decreased expression of the fibrosis-associated gene *Col1a1*. These results suggest that inhibition of TGFβ/SMAD signaling is an important mechanism underlying the anti-fibrotic effects of IVA337.

Inflammatory activation is a major contributor to lymphatic remodeling and fibrosis. We found that IVA337 significantly reduced inflammatory cell infiltration in lymphedematous skin, consistent with the known anti-inflammatory properties of PPAR activation. PPAR signaling can suppress inflammatory pathways, including NF-κB activation, and reduce the expression of pro-inflammatory mediators ^[32]^. In addition, targeting CD4^⁺^ T cell-mediated inflammation has been shown to ameliorate lymphedema in experimental models, highlighting the critical role of adaptive immunity in disease progression ^[33,34]^. Given the well-established immunomodulatory effects of PPAR activation ^[21]^, it will be important to determine whether IVA337 influences the expression of chemokines and cytokines, as well as the recruitment, activation, and polarization of T cells during lymphedema progression. Future work elucidating these mechanisms may further clarify the immunoregulatory actions of IVA337 and support its therapeutic potential for lymphedema.

In addition to chronic inflammation and fibrosis, impaired cholesterol clearance has recently been recognized as a pathogenic mechanism contributing to lymphedema progression ^[35]^. Notably, IVA337 also regulates the lipid metabolism through the activation of PPARs, making it interesting to determine whether IVA337 also improves cholesterol clearance during lymphedema progression.

Although the anti-inflammatory and anti-fibrotic properties of PPARα, PPARβ/δ, and PPARγ have been well documented ^[12,19,20]^, the relative contribution of each isoform to the therapeutic effects of IVA337 in lymphedema remains unclear. Future studies using isoform-specific agonists, antagonists, or genetic models will be necessary to define the mechanisms underlying pan-PPAR activation. In addition, our study focused on the early stage of secondary lymphedema; therefore, the long-term efficacy of IVA337 in established fibrotic lymphedema warrants further investigation.

In summary, our study reveals an unexpected role of IVA337 in lymphatic disease and demonstrates that pan-PPAR activation alleviates secondary lymphedema by suppressing TGFβ/SMAD2/3-mediated fibrotic remodeling, reducing inflammation, and preserving lymphatic endothelial integrity. These findings provide a mechanistic foundation for developing PPAR-targeted therapies to prevent lymphatic dysfunction and progressive fibrosis in secondary lymphedema.

## Author Contributions

Conceptualization, J.P. and X.L.; study design, J.P. and X.L.; performed experiments, J.P.; mouse experiments and tissue analysis, J.P.; imaging and quantitative analysis, J.P.; molecular experiments, J.P.; data analysis and interpretation, J.P. and X.L.; writing—original draft preparation, J.P.; writing—review and editing, J.P., L.D., E.D, L.F., J.Z., and X.L.; supervision, X.L.; funding acquisition, X.L.. All authors have read and agreed to the published version of the manuscript.

## Acknowledgements

We thank Lemole Center Imaging Core for providing advanced microscope and equipment support in this study. Graphic image was created in BioRender. Do, L. (2026) https://BioRender.com/se09h7e.

## Source of funding

This work was supported by NIH grant 1RO1HL163269, DOD grants HT942524PRMRPDA, HT9425-25-1-0034 to X.L., American heart association predoctoral fellowship grant 26PRE1561542 to E.D..

## Disclosure

None

## Abbreviations

PPAR: peroxisome proliferator-activated receptor
MASH: metabolic dysfunction-associated steatohepatitis;
TGFβ: transforming growth factor beta
SMAD: sma- and mad-related proteins
HDLECs: human dermal lymphatic endothelial cells
Cpt1a: carnitine palmitoyltransferase 1a
Fabp4: fatty acid binding protein 4
CDH5: cadherin 5
VE-Cadherin: vascular endothelial cadherin
TJP1: tight junction protein 1
CLDN5: claudin5.

## Notes

https://BioRender.com/se09h7e

### Competing Interest Statement

The authors have declared no competing interest.

