## supplementary file for "Pan-Peroxisome Proliferator-Activated Receptor Agonist IVA337 Alleviates Secondary Lymphedema via Inhibiting TGFβ/SMAD2/3 Signaling Pathway"

### S-Figure 1

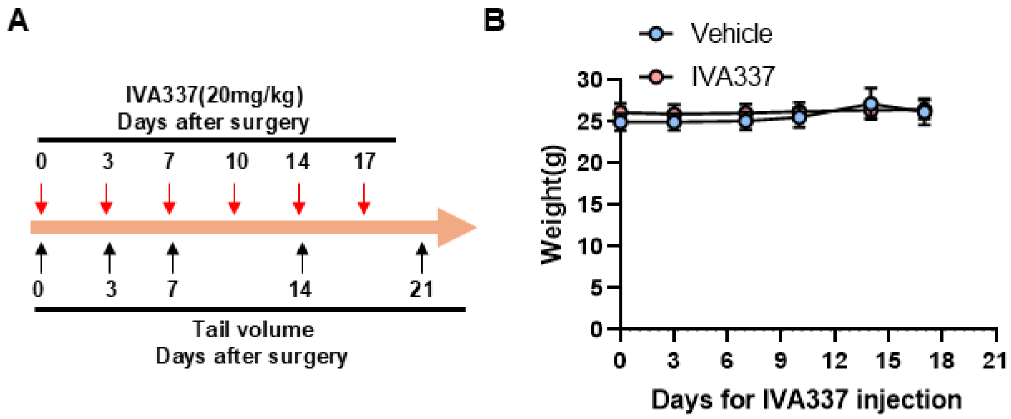


**Supplemental Figure 1. Mouse wight was stable in the process of IVA337 administration. A.** The timeline of tail surgery and IVA337 administration. **B.** The mouse weight was quantified and analyzed in the process of IVA337 injection. N=4-6.

### S-Figure 2

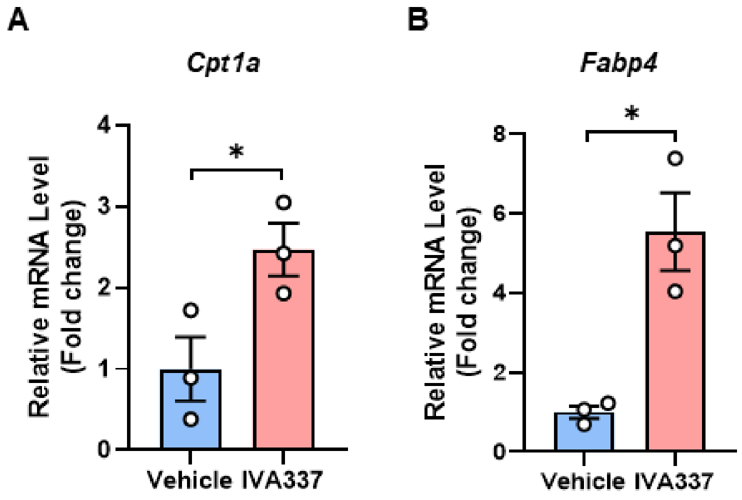


**Supplemental Figure 2. IVA337 activates PPARs. A-B.** Mouse tail bulk-RNA was isolated, and qRT-PCR was used to check the expression of *Cpt1a*, and *Fabp4*, which are classic PPAR target genes. N=3. Data are shown as mean ± SEM, *P<0.05, unpaired two-tailed Student's t-test.

S-Figure 3

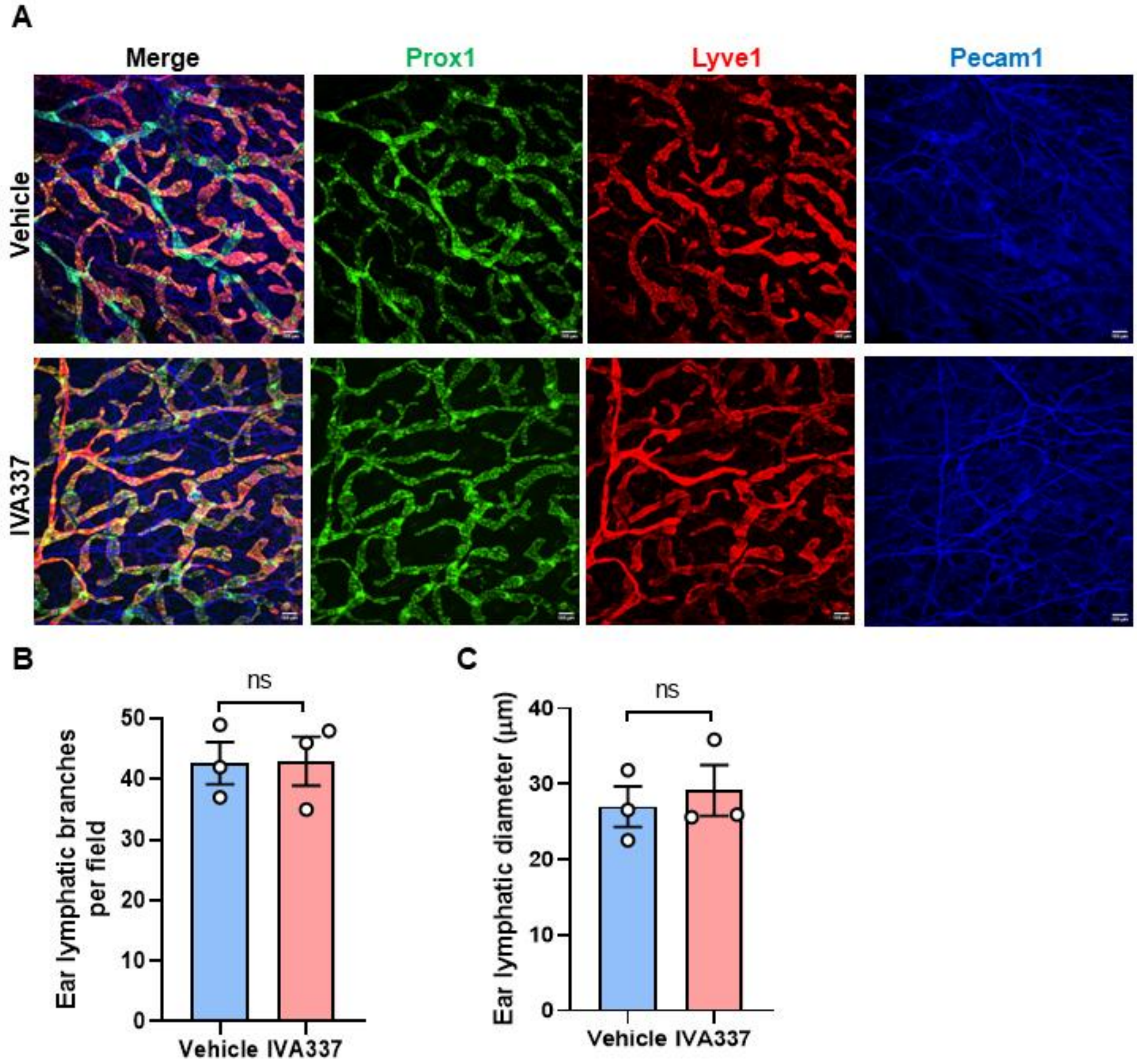


**Supplemental Figure 3. Ear lymphatic vessels were not influenced by IVA337.**

**A-C.** Ear whole mount staining targeted Lyve1, Prox1, and Pecam1, lymphatic vessels numbers and diameter were quantified and analyzed. N=3. Scale bar: 100 μm, Data are shown as mean ± SEM, unpaired two-tailed Student's t-test.

S-Figure 4

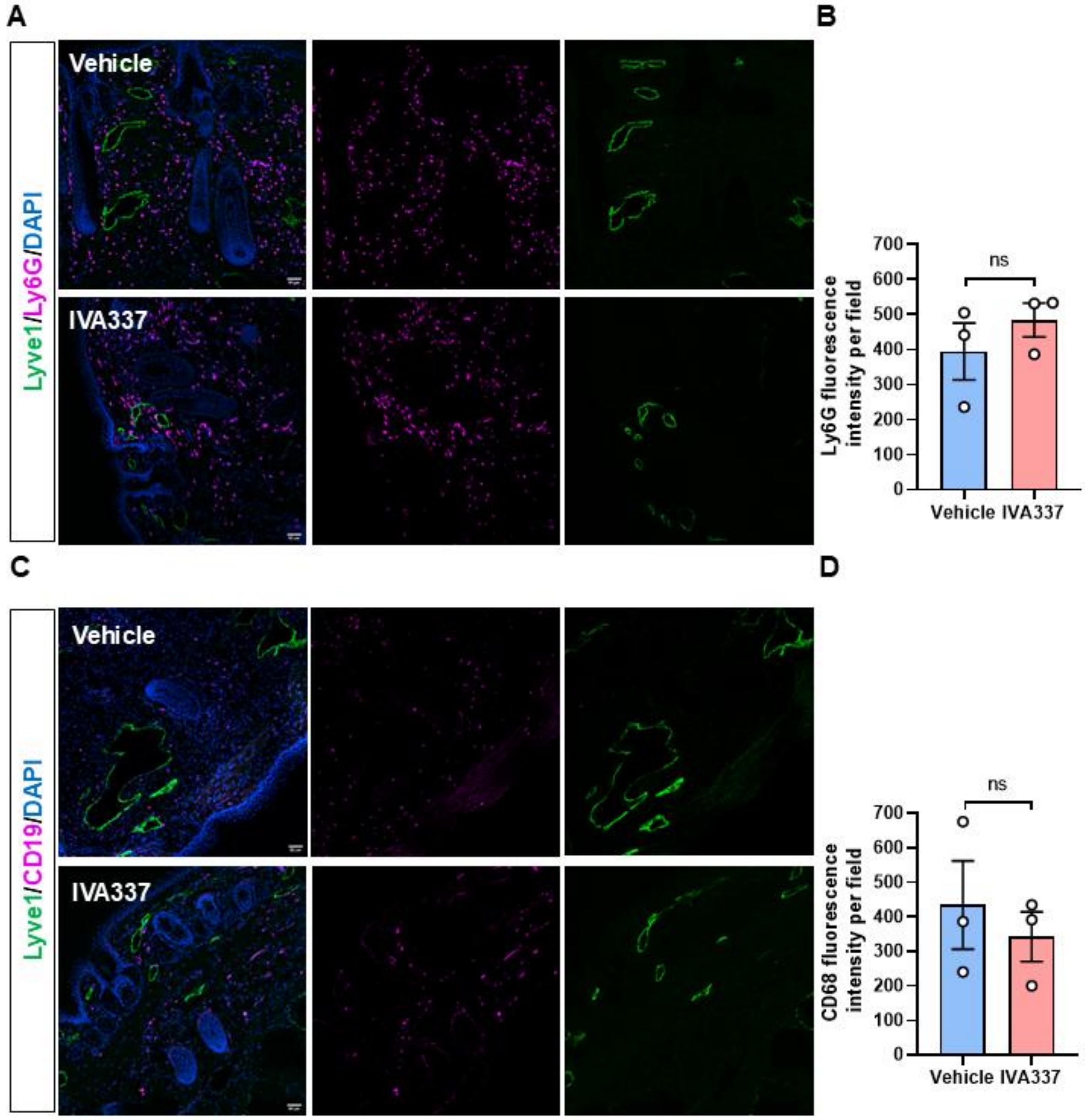


**Supplemental Figure 4. Neutrophils and B cells were not changed between control and IVA337 groups. A-B.** Immunostaining targeted for neutrophils marker Ly6G together with LECs marker Lyve1 in tail cryosection, and quantification and analysis of fluorescent intensity in vehicle and IVA337 group. N=3. **C-D.** Immunostaining targeted for B cells marker CD19 together with LECs marker Lyve1 in tail cryosection, and quantification and analysis of fluorescent intensity in vehicle and IVA337 group. N=3. Scale bar is 50 μm. Data are shown as mean ± SEM, unpaired two-tailed Student's t-test.

S-Figure 5

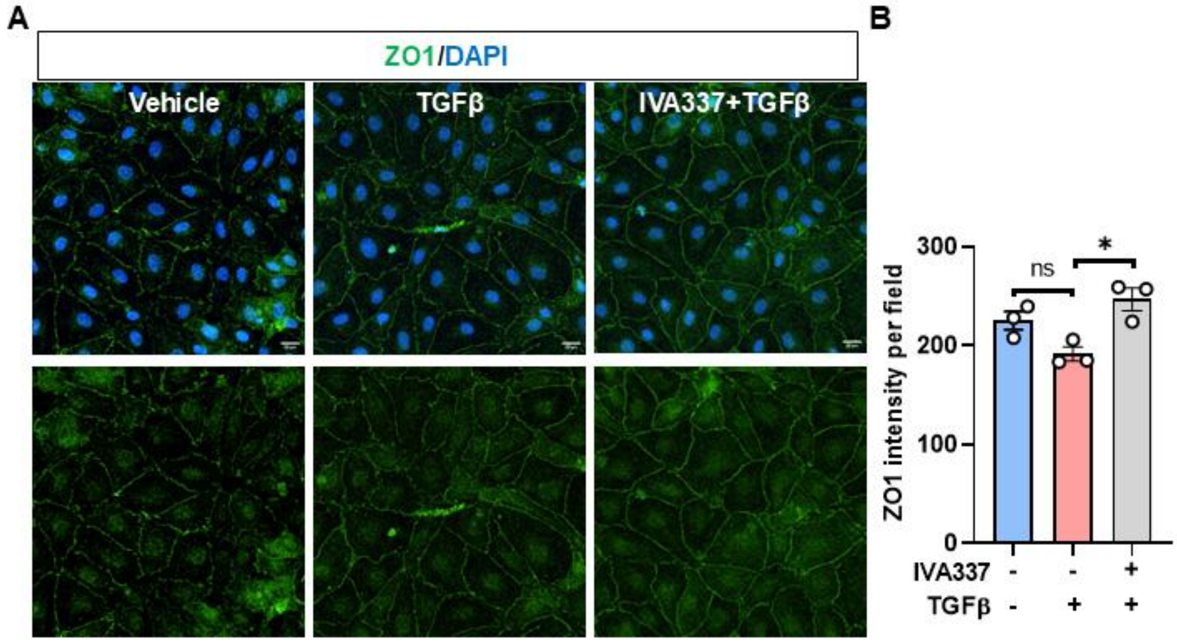


**Supplemental Figure 5. IVA337 rescued TGFβ-induced ZO1 expression in HDLECs. A-B.** HDLECs were pretreated IVA337 (5 μM, 16h) and then treated with TGFβ (20 ng/ml, 8 h), immunostaining targeted for ZO1 and DAPI to check the cell junction. Fluorescent intensity was analyzed in B. N=3. Scale bar is 20 μm. Data are shown as mean ± SEM, *P<0.05, one-way ANOVA followed by Turkey's test.

S-Figure 6

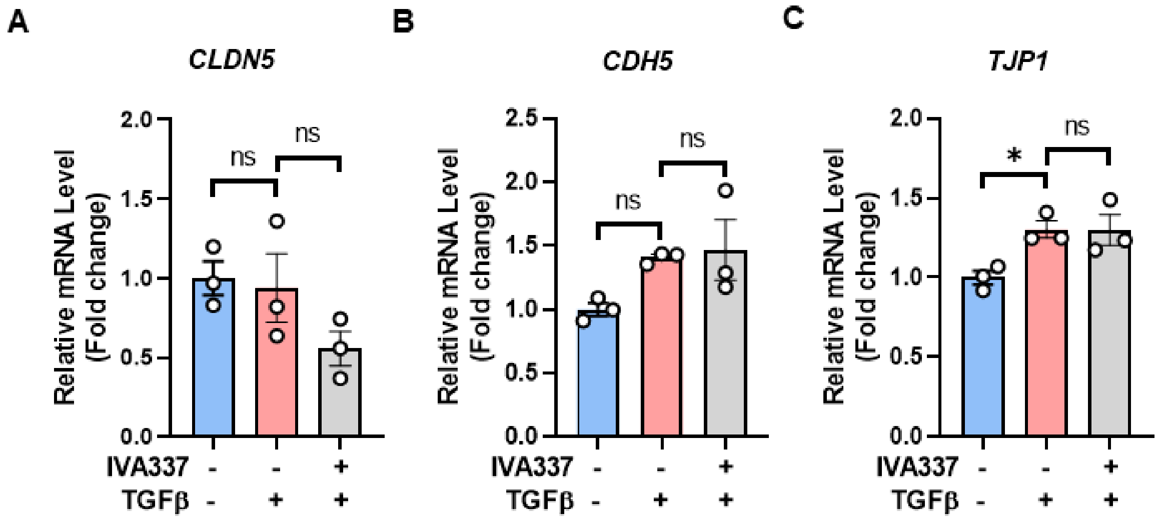


**Supplemental Figure 6. IVA337 has no effects on the mRNA levels of *CLDN5*, *CDH5*, and *TJP1* in HDLECs. A-C.** HDLECs were pretreated IVA337 (5 μM, 16h) and then treated with TGFβ (20 ng/ml, 8 h), RNA was isolated and RT-PCR was used to check the expression of *CLDN5*, *CDH5*, and *TJP1*. Data are shown as mean ± SEM, *P<0.05, one-way ANOVA followed by Turkey's test.

Supplementary Table S1.

| Target antigen | Vendor | Catalog Number | Working Concentration |
|---|---|---|---|
| Prox1 | Abcam | ab101851 | IF 1:500 |
| Pecam1 | BD Bio | 557355 | IF 1:500 |
| VE-Cadherin | R and D systems | AF1002 | IF 1:1000 |
| GAPDH | ThermoFisher | MA5-15738-HRP | WB 1:5000 |
| ZO1 | Invitrogen | 61-7300 | IF 1:500 |
| Claudin5 | Invitrogen | MA5-32614 | IF 1:500 |
| p-SMAD | Abcam | AB52903 | IF 1:500; WB 1:1000 |
| t-SMAD | CST | 3102 | WB 1:1000 |
| CD68 | Abcam | AB125212 | IF 1:500 |
| CD45 | BD Biosciences | 553076 | IF 1:500 |
| CD8a | BD Biosciences | 553028 | IF 1:500 |
| Ly6G | Biolegend | 127602 | IF 1:500 |
| CD19 | ThermoFisher | 14-0194-82 | IF 1:500 |
| Lyve1 | Invitrogen | 14044382 | IF 1:1000 |
| Donkey anti-rat, Alexa Fluor™ 488 | ThermoFisher | A-21208 | IF 1:1000 |
| Donkey anti-goat, Alexa Fluor™ 488 | ThermoFisher | A-11055 | IF 1:1000 |
| Donkey anti-goat, Alexa Fluor™ 594 | ThermoFisher | A-11058 | IF 1:1000 |
| Donkey anti-rabbit, Alexa Fluor™ 647 | Thermofisher | A-31573 | IF 1:1000 |
| Donkey anti-Rabbit Alexa Fluor™ 594 | Invitrogen | A-21207 | IF 1:1000 |
| Donkey anti-goat, Alexa Fluor™ 647 | Invitrogen | A-21447 | IF 1:1000 |
| CY5 donkey anti-Rabbit | Jackson Immuno | 711-175-152 | IF 1:1000 |
| CY3 donkey anti Rat | Jackson Immuno | 112-165-003 | IF 1:1000 |
| Cy5 donkey anti-Mouse | Jackson Immuno | 715-175-150 | IF 1:1000 |
| CY5 donkey anti-Rat | Jackson Immuno | 712-175-150 | IF 1:1000 |

Supplementary Table S2

| Gene | Species | Forward (5’-3’) | Reverse (5’-3’) |
|---|---|---|---|
| *GAPDH* | Human | GGTGTGAACCATGAGAAGTATGA | GAGTCCTTCCACGATACCAAAG |
| *CLDN5* | Human | TGCAAGAGTGTGCTAGAGGC | ACAAAGCAGTCCACGAGGTAG |
| *CDH5* | Human | GAAGCCTCTGATTGGCACAGTG | TTTTGTGACTCGGAAGAACTGGC |
| *TJP1* | Human | GTCCAGAATCTCGGAAAAGTGCC | CTTTCAGCGCACCATACCAACC |
| *18S* | Mouse | GAAACGGCTACCACATCCAAGG | GCCCTCCAATGGATCCTCGTTA |
| *Col1a1* | Mouse | GCTCCTCTTAGGGGCCACT | CCACGTCTCACCATTGGGG |

| | | | |
|---|---|---|---|
| *Cpt1a* | Mouse | TGGCATCATCACTGGTGTGTT | GTCTAGGGTCCGATTGATCTTTG |
| *Fabp4* | Mouse | AAGGTGAAGAGCATCATAACCCT | TCACGCCTTTCATAACACATTCC |
